# Identification of cellular double-stranded RNAs in mammalian embryonic stem cells

**DOI:** 10.64898/2026.04.13.718158

**Authors:** Katharina J. Kases, Pilar G. Marchante, Jeroen Witteveldt, Guillermo Peris, Sara R. Heras, Sara Macias

## Abstract

Antiviral defence mechanisms are typically activated upon sensing virus-derived nucleic acids. During replication, viruses generate double-stranded RNA (dsRNA) intermediates that the innate immune system can sense, triggering several defence pathways. Conversely, mammalian cells avoid accumulating their own endogenous dsRNA to prevent activating these defence mechanisms. However, we demonstrate that mammalian embryonic stem cells (ESCs) accumulate endogenous dsRNA without activating these responses, as they lack all classical dsRNA-mediated antiviral pathways. To identify these endogenous dsRNAs, we have developed a method that includes an antibody-based purification coupled to RNase I treatment to enrich bona fide dsRNAs. The RNase I treatment results in an enrichment on sense/antisense and A-to-I edited transcripts, suggesting that they are true dsRNA in cells. Our refined protocol reveals that transposable elements (TEs), primarily the young elements from the LINE and LTR classes, are the predominant sources of dsRNA in ESCs. This approach will be useful for investigating the role of dsRNA in disease settings, such as autoimmunity or cancer, where endogenous dsRNA accumulation has also been observed.

## Introduction

The accumulation of virus-derived double stranded RNA (dsRNA) in the cytoplasm of mammalian cells triggers multiple defence pathways. Viral dsRNA is sensed by the retinoic acid-inducible gene I (RIG-I)-like receptor (RLR) family of proteins, which activates the major antiviral innate immune pathway, the type I interferon (IFN) response. Within the RLR family, RIG-I recognises short and uncapped dsRNAs, while MDA5 recognises longer dsRNAs (Rehwinkel and Gack 2020). Upon dsRNA recognition, both sensors activate the transcription factor IRF3 driving the expression of type I interferons (IFNs) (Bourdon et al. 2025). Within the endosomal compartment, the Toll-like receptor 3 (TLR3) sensor also recognises dsRNA leading to IFN expression through IRF3 activation (Chattopadhyay and Sen 2014). In addition to IFNs, viral dsRNAs can induce other antiviral responses. For instance, upon binding to dsRNA, the protein kinase R (PKR) phosphorylates the translation initiation factor eIF2α, causing a general inhibition of cap-dependent mRNA translation (Dauber and Wolff 2009). DsRNA also activates the OAS/RNase L pathway resulting in a global degradation of both cellular and viral RNA (Lee et al. 2026). Although these dsRNA-activated responses are critical in providing antiviral defence, they can also be deleterious if inappropriately activated.

Considering that the innate immune system cannot discriminate virus from host-derived dsRNA molecules, mammalian cells avoid accumulating endogenous dsRNAs to prevent aberrant immune activation. Mammalian cells possess several mechanisms to prevent dsRNA accumulation, one of the most studied being based on RNA modifications. The post-transcriptional editing of adenosine to inosines (A-to-I) within dsRNA by ADAR disrupts base-pairing, preventing their sensing by MDA5 or PKR (Rehwinkel and Mehdipour 2025). Most editing sites are contained in the non-coding regions of RNAs, overlapping with repetitive sequences derived from transposable elements (TEs), predominantly SINE/Alu elements (Porath et al. 2017). M6A modifications have also been implicated in suppressing the RLR- dependent IFN response, as loss of the m6A writer METTL3 resulted in an increased IFN signature, suggesting that m6A modifications prevents dsRNA formation (Gao et al. 2020). Similarly, incorporation of pseudouridines into mRNA prevents sensing by PKR (Anderson et al. 2010). Additional mechanisms include the activity of RNA helicases, such as DHX9, which has been proposed to inhibit the formation of TE-derived dsRNAs in complex with other dsRNA binding proteins, including ADAR1 or DGCR8 (Aktaş et al. 2017; Murayama et al. 2024; Gázquez-Gutiérrez et al. 2026). Finally, dsRNA structures within non-coding regions of mRNAs can be resolved through Staufen-mediated mRNA decay triggered by dsRNAs in the 3′UTR of protein-coding transcripts (Gong et al. 2013). More recently, a distinct splicing pathway, termed SOS splicing, was shown to remove inverted terminal repeat elements adopting dsRNA hairpin structures, thereby restoring the function of genes disrupted by TEs (Zhao et al. 2026).

The repetitive nature of TEs, along with the fact that they can be transcribed in sense or antisense orientation as part of the genes where they are inserted, makes them prime candidates for dsRNA formation. As such, TEs have emerged as a major source for dsRNA formation in different disease settings, including cancer, senescence and autoimmune diseases (Gázquez-Gutiérrez et al. 2021). Transcription initiated from intrinsic TE promoters is epigenetically silenced across most healthy somatic tissues to minimise the risk of de novo transposition and consequent genomic instability. A notable exception occurs during early development, when specific TE families are derepressed and can be transcribed as full-length copies from their own promoters. Such is the importance of TE expression in embryogenesis, that they orchestrate the transcriptional profile during early development (DiRusso and Clark 2023; Oomen et al. 2025).

To identify mammalian cellular dsRNAs, multiple methods have been developed. DsRNAs can be captured using antibodies against dsRNA with either purified RNA or total cell extracts as input material (Dhir et al. 2018; Wiatrek et al. 2019; Gao et al. 2020; Ghosh et al. 2020; Krieger et al. 2024; Witteveldt et al. 2025; Gázquez-Gutiérrez et al. 2026). One of the most well-studied antibodies is the monoclonal J2, which is highly specific against dsRNA, and does not show cross-reactivity against dsDNA, RNA/DNA hybrids, ssRNA, or ssDNA. J2 was initially reported to recognise dsRNA species with a minimum length of 40 base pairs (bp); however, more sensitive recent studies indicate that it can bind dsRNA helices as short as 14 bp, with binding affinity increasing with dsRNA length (Schonborn et al. 1991; Bou-Nader et al. 2025). Besides antibodies, dsRNA-binding proteins, such as dsRNA sensors, can also be used to identify cellular dsRNAs. For instance, *in vitro* RNase protection assays with MDA5 and immunoprecipitation of MDA5 have been used to identify cellular dsRNAs that can bind this sensor (Ahmad et al. 2018; Clapes et al. 2021). Similarly, PKR immunoprecipitation following formaldehyde cross-linking has been successfully used to identify cellular dsRNAs (Kim et al. 2018). Nonetheless, using sensors for dsRNA identification has limitations, as they only capture the subset of transcripts preferentially bound by a given sensor, rather than the entire population of dsRNAs in the cell.

Here, we have developed a method to identify the global landscape of bona fide dsRNAs using the anti-dsRNA antibody J2 combined with RNase I digestion. As a model system, we used embryonic stem cells (ESCs) which provide a permissive environment for dsRNA accumulation. Our recent work has shown that, although mouse ESCs accumulate significantly higher levels of dsRNA than somatic cells, they fail to elicit a type I interferon (IFN) response (Witteveldt et al. 2025). We now show that in addition to the IFN response, mouse ESCs lack all major antiviral responses to dsRNA, including PKR and the OAS/RNase L pathways, and are therefore unable to respond to either exogenous or endogenous dsRNA. Our immunoprecipitation strategy exploits the differential sensitivity of dsRNA to structure-specific RNA nucleases, including RNase I and RNase III. Whereas dsRNA is largely resistant to single-strand-specific nucleases such as RNase I, it remains sensitive to cleavage by the dsRNA-specific endoribonuclease RNase III. RNase I treatment coupled with J2 immunoprecipitation has been successfully used to identify virus-derived dsRNAs (Decker et al. 2019; Price et al. 2022). Our results demonstrate that RNase I digestion enhances the detection of bona fide cellular dsRNAs by enriching for both sense, antisense and edited transcripts, while removing non-dsRNA sequences. We further show that dsRNAs in ESCs derive from TEs, and are enriched for the youngest LINE and LTR subfamilies. These evolutionary young elements are typically longer and accumulate fewer mutations, features that increase their propensity to form highly complementary and therefore more stable dsRNAs. This approach should improve both the sensitivity and specificity of bona fide dsRNA identification across diverse biological contexts, including homeostasis, infection, and pathological states associated with aberrant dsRNA accumulation, such as autoimmunity and cancer.

## Results

### mESCs lack dsRNA-mediate immune responses

We and others have previously shown that mouse ESCs cannot trigger the IFN response upon dsRNA stimulation (Wang et al. 2013; Witteveldt et al. 2019, 2025). By contrast, the status of other major dsRNA-induced innate immune pathways in mouse ESCs remains poorly defined. To address this, we transfected mESCs with exogenous dsRNA (poly(I:C)) and compared their response with that of their differentiated derivatives. As expected, ESC differentiation resulted in decreased expression of the pluripotency markers *Nanog* and *Oct4* (*Pou5f1*) and increased expression of the differentiation markers *Acta2* and *Tagln*, as shown by RT-qPCR. None of these markers were significantly affected by the dsRNA challenge (**Figure 1a**). In differentiated cells, dsRNA stimulation activated the expression of IFNs (*Ifnb1*) and interferon-stimulated genes (ISGs, *Oas1* and *Stat1*), while no increase in expression was observed in ESCs, as expected (**Figure 1b**). In agreement with these findings, differentiated cells exhibited phosphorylated IRF3 upon dsRNA stimulation, indicating successful RLR-mediated signalling (**Figure 1c** and **Supplementary Figure 1**). Next, we assessed whether ESCs preserved other dsRNA-mediated antiviral responses, specifically the PKR and OAS/RNase L pathways. We monitored the phosphorylation of the kinase PKR by western blot, as this kinase auto-phosphorylates upon dsRNA binding. Interestingly, PKR phosphorylation was detected only in differentiated cells, with no changes observed in ESCs upon dsRNA transfection (**Figure 1d** and **Suppl Fig 1**). Finally, dsRNA stimulation also activated the OAS/RNase L system in differentiated cells, leading to degradation of ribosomal RNA (rRNA). In contrast, ESCs did not respond to dsRNA, maintaining the integrity of both 28S and 18S rRNA (**Figure 1e**). These findings indicate that all the major dsRNA-mediated antiviral responses are inactive in ESCs and are only acquired upon differentiation. This led us to conclude that mESCs represent a dsRNA-permissive environment.

**Figure 1.**
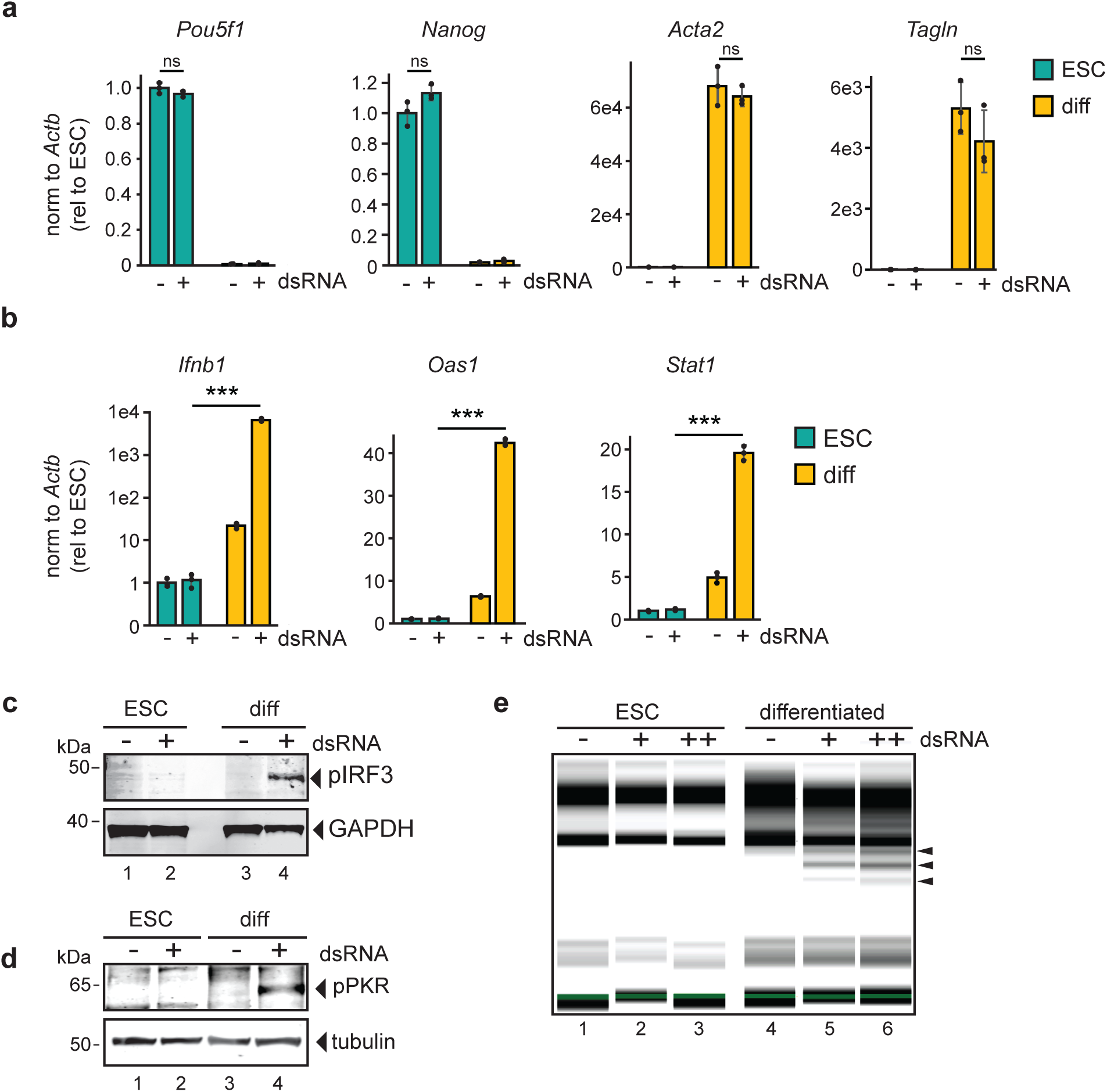
Characterisation of dsRNA-mediated antiviral response in ESCs and differentiated cells. (*A*) Expression of the pluripotency markers *Pou5f1* and *Nanog* and differentiation markers *Acta2* and *Tagln* in ESCs and differentiated cells was measure by RT-qPCR, comparing mock and dsRNA-stimulated cells. *Actb* was used as a normaliser. Data represent the average of three biological replicates ± SD relative to untreated ESC. One-way ANOVA was used to calculate significant differences between dsRNA- treated samples within each cell line, followed by a Tukey HSD. *p-val ≤ 0.05, **p-val ≤ 0.01, ***p-val ≤ 0.001. (*B*) Expression of *Ifnb1* and interferon stimulated genes *Oas1* and *Stat1* in ESCs and differentiated cells upon dsRNA or mock stimulation by RT-qPCR analysis. *Actb* was used as a normaliser. Data represent the average of three biological replicates ± SD relative to untreated ESC. One-way ANOVA was used to calculate significant differences between dsRNA-treated cell lines, followed by a Tukey HSD. *p-val ≤ 0.05, **p-val ≤ 0.01, ***p-val ≤ 0.001. (*C*) Western blot analysis of phosphorylated IRF3 levels in ESCs and differentiated cells. GAPDH serves as a loading control (*D*) Western blot analysis of phosphorylated PKR in ESCs and differentiated cells. Tubulin serves as a loading control (*E*) rRNA electrophoresis from ESCs and differentiated cells, mock and dsRNA-stimulated. (+) represents lower dsRNA and (++) higher dsRNA stimulation.

### Accumulation and purification of dsRNAs in ESCs

Compared with somatic cells, mESCs accumulate endogenous dsRNA (Witteveldt 2025). However, because ESCs lack the major antiviral pathways triggered by dsRNA, this accumulation does not trigger any unwanted immune activation. By contrast, these antiviral pathways become activate upon differentiation. This observation led us to hypothesise that endogenous dsRNA levels must be reduced during differentiation to prevent its aberrant recognition as foreign. To test this, we compared the levels of dsRNA in ESCs versus *in vitro* differentiated cells using anti dsRNA antibodies (J2) by flow cytometry. Our findings confirmed that ESCs but not differentiated cells accumulate endogenous dsRNAs (**Figure 2a-b**). We verified the specificity of the dsRNA signal by treating cells with RNase III, a dsRNA- specific nuclease, before staining. RNase III treatment resulted in a reduction of the signal, approaching the level found in differentiated cells (**Figure 2a-b, Supplementary Figure 2**). Immunofluorescence with the dsRNA antibody confirmed that ESCs accumulate substantial amounts of dsRNA and that this accumulation is lost upon differentiation (**Figure 2c**). These results demonstrate that endogenous dsRNAs accumulate in ESCs but are lost upon differentiation, coincident with the acquisition of functional dsRNA-mediated responses.

**Figure 2.**
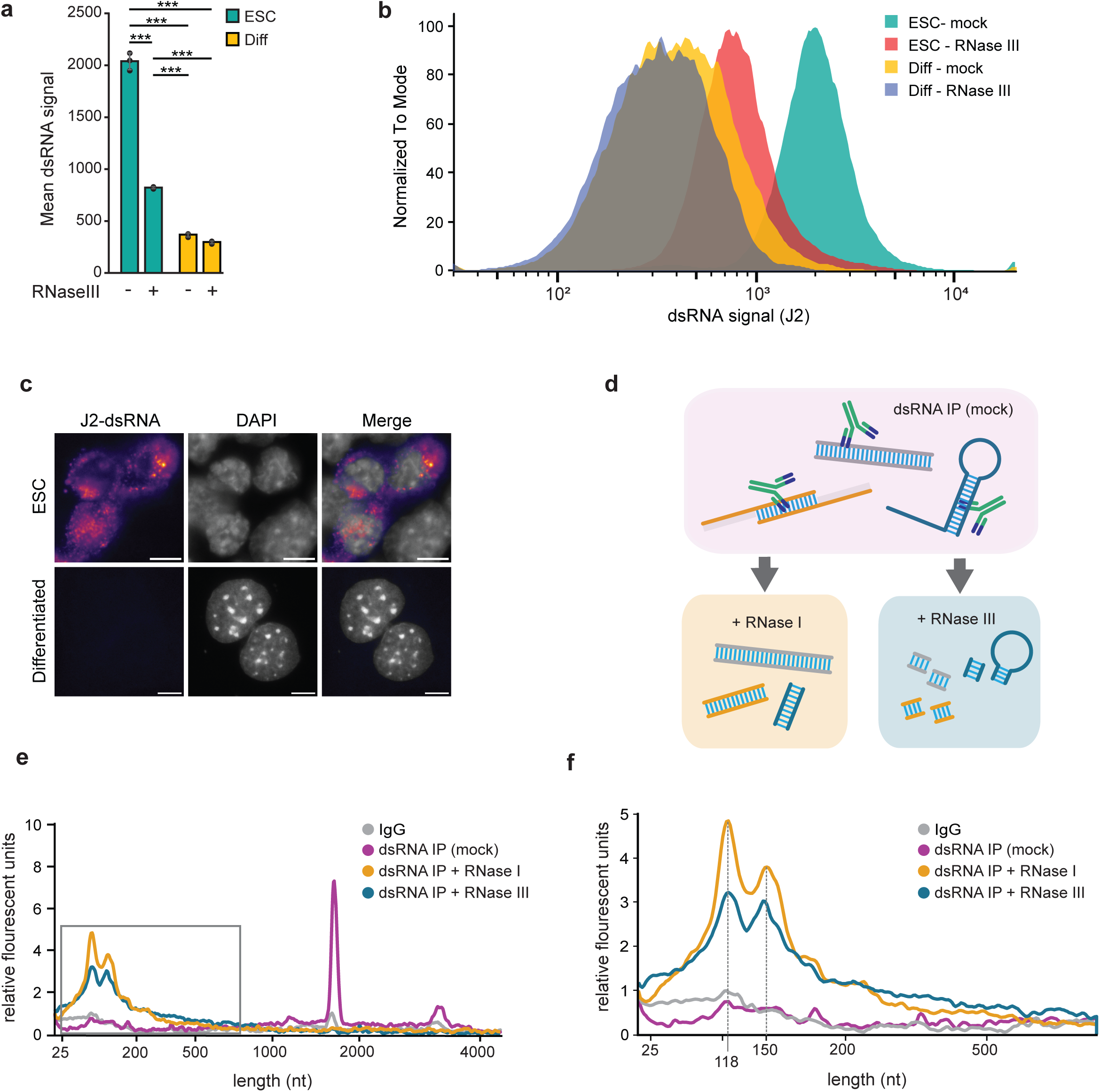
DsRNA quantification in ES and differentiated cells. (*A*) Flow cytometry analysis against dsRNA using the dsRNA-specific antibody J2 in ESC vs differentiated (diff) cells in the presence or absence of RNase III. RNase III treatment tests specificity of J2 antibody signal. Data represent the average of three biological replicates ± SD. One-way ANOVA was used to calculate significant differences amongst comparisons, followed by a Tukey HSD. *p-val ≤ 0.05, **p-val ≤ 0.01, ***p-val ≤ 0.001. (*B*) Representative examples of normalised Flow cytometry intensity in ESCs and differentiated cells after mock and RNaseIII treatment. For all replicates and gating strategy see **Supplementary figure 2a** (*C*) Immunofluorescence for dsRNA. J2 antibody was used to detect dsRNA, and DAPI to stain the nucleus. Scalebar is 10 µm (*D*) Setup of dsRNA-immunoprecipitation using the J2 antibody. DsRNAs bound to J2 were either mock treated, treated with RNaseI or RNaseIII prior to purification and followed high-throughput RNA sequencing. (*E*) Bioanalyzer electropherogram of purified RNAs J2-IP using the different treatments shown in (*D*). (*F*) Detailed view of Bioanalyzer electropherogram showing accumulation of shorter fragments upon treatment with RNase I and III.

To identify the ESC-enriched dsRNAs, we developed an improved J2-based immunoprecipitation protocol, coupling immunoprecipitation with RNase I digestion to remove single-stranded RNA (ssRNA) regions. RNAse I is an endoribonuclease that specifically degrades ssRNA without sequence bias (Kennell 2002). In cells, dsRNAs can arise by complementary interactions either *in cis* within a single transcript or *in trans* between distinct RNA molecules. Intramolecular base-pairing of complementary sequences can give rise to hairpin structures, whereas intermolecular pairing between sense and antisense transcripts can generate dsRNA duplexes. These structures may be perfectly base-paired or may contain mismatches resulting in bulges, internal loops and single-stranded overhangs. (**Figure 2d**, top). Given that the J2 antibody requires only a minimal dsRNA stretch for binding, we hypothesised that besides this minimal duplex region, many of the immunoprecipitated RNAs would still retain substantial single stranded regions. To enrich for bona fide dsRNA species in our samples, we treated J2-immunoprecipitates with RNase I while still bound to the J2 antibody (**Figure 2d**, bottom left). In parallel, to confirm that the immunoprecipitated RNAs represented bona fide dsRNAs, we treated the J2-immunoprecipitated samples with RNAse III, a dsRNA-specific endonuclease (**Figure 2d**, bottom right). The RNA recovered after immunoprecipitation and treatment with RNase I, RNAse III or no-treatment (‘mock’) was analysed on a Bioanalyzer alongside an IgG control. This confirmed the specificity of the J2 immunoprecipitation, as no RNA was recovered from the IgG control (**Figure 2e**). Compared with the mock-treated dsRNA IP, both RNase treatments reduced the size of the predominant immunoprecipitated RNA species (**Figure 2f**). Overall, these RNA profiles demonstrated that the J2 immunoprecipitation was specific, and that treatment with either nuclease enriched for smaller RNA fragments.

### RNase I treatment enhances the detection of bona fide dsRNAs

Immunopurified dsRNAs were subsequently identified using high-throughput RNA sequencing. Since the IgG negative control did not yield sufficient RNA for library preparation, we used total RNA-seq of the input samples to calculate the enrichment of immunoprecipitated dsRNAs. Principal component analysis revealed that all the immunoprecipitated dsRNA samples clustered into distinct groups and, regardless of RNase treatment, were enriched for repetitive elements, including transposable elements (TEs), relative to the input (**Supplementary Figure 3a-b**). Differential enrichment analyses showed an enrichment across all classes of transposable elements following dsRNA immunoprecipitation (**Figure 3a**). However, these TE classes exhibited different sensitivities to RNase treatments. All classes were sensitive to RNase III relative to the untreated dsRNA immunoprecipitations (‘mock’), confirming their dsRNA content. However, unlike other TE classes, LINEs were similarly enriched upon RNase I treatment, suggesting that LINE-derived dsRNA species have the highest levels of complementarity, making them more resistant to RNase I degradation (**Figure 3a**).

**Figure 3.**
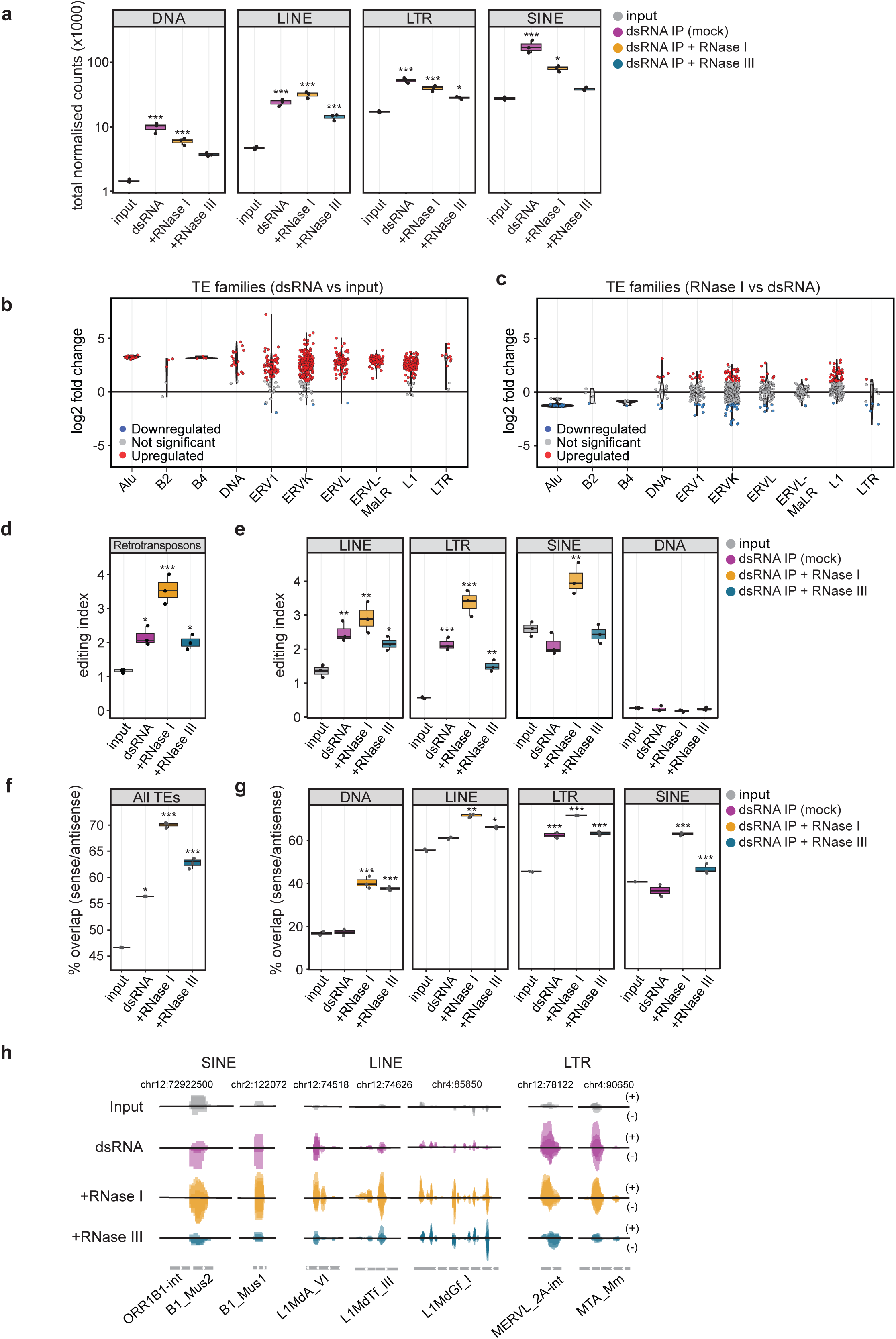
DsRNAs are enriched in transposable elements with RNAse treatment-specific differences. (*A*) Normalised counts of the four major classes of transposable elements compared to input controls. One-way ANOVA was used to calculate significant differences between input and immunoprecipitated samples, followed by a Tukey HSD. *p-val ≤ 0.05, **p-val ≤ 0.01, ***p-val ≤ 0.001. (*B*) Violin plots showing differential enrichment (log₂ fold change) of TE subfamilies within selected families for dsRNA IP (mock) versus input. Points represent individual subfamilies, coloured by differential enrichment (red: upregulated, blue: downregulated, grey: not significant; log₂FC > 1, adjusted p < 0.05). (*C*) Violin plots showing differential enrichment (log₂ fold change) of TE subfamilies within selected families after RNase I treatment versus dsRNA (mock). Points represent individual subfamilies, coloured by differential enrichment (red: upregulated, blue: downregulated, grey: not significant; log₂FC > 1, adjusted p < 0.05). (*D*) Levels of A-to-I editing in retrotransposons for input and dsRNA-IPs. One-way ANOVA was used to calculate significant differences between input and immunoprecipitated samples, followed by a Tukey HSD. *p-val ≤ 0.05, **p-val ≤ 0.01, ***p-val ≤ 0.001. (*E*) Levels of A-to-I editing by TE class. Statistical analysis as in (*D*). (*F*) Percentage of sense reads mapping to TEs with a corresponding antisense match, for input and dsRNA IPs. One-way ANOVA was used to calculate significant differences between input and immunoprecipitated samples, followed by a Tukey HSD. *p-val ≤ 0.05, **p-val ≤ 0.01, ***p- val ≤ 0.001. (*G*) Sense/antisense overlap by TE class. Statistical analysis as in (*F*). (*H*) Examples of specific TE locus. The combined normalised reads from the biological replicates mapped are shown, for inputs and dsRNA IPs. Above black line represents sense (or ‘plus’), below represents antisense reads (or ‘minus’). Chromosomal location of each locus is represented at the top

DsRNA IPs also displayed significant enrichment for most TE families compared to input (**Figure 3b**). They were enriched in SINEs, including Alu, B2 and B4 families; the LTR families ERV1, ERVK, ERVL, ERVL-MaLR and LTR in addition to DNA and LINE families (**Figure 3b**). In agreement with our previous analysis, RNase I treatment resulted in a reduction in the Alu, B2 and B4 families. A less consistent behaviour was observed for LTR families, as some were enriched or lost upon RNAse I treatment. In contrast, a high proportion of LINE subfamilies were enriched upon RNAse I addition, suggesting that LINEs form highly complementary dsRNAs, while the others may be more frequently interrupted by ssRNA regions (**Figure 3c**). To further confirm that the TE families enriched after RNase I represent bona fide dsRNA species, we assessed their behaviour after RNase III treatment. Most TE subfamilies showed significant depletion following dsRNA cleavage, supporting the conclusion that RNase I treatment preferentially enriches for bona fide dsRNA species (**Supplementary figure 4a**).

To verify that the enriched sequences derived from bona fide dsRNAs, we also quantified adenosine-to-inosine (A-to-I) editing levels, a marker of ADAR activity on dsRNA. The presence of editing also indicates that these dsRNAs were present in cells, rather than being interactions formed in extracts or during cell lysis. Across all retrotransposons, RNase I-treated samples showed the highest editing levels, further supporting the conclusion that RNase I treatment enriches bona fide dsRNA species (**Figure 3d**). When differentiating between TE classes, we observed similar patterns across the board with the highest editing levels for SINE/Alu elements after RNAse I treatment, which are considered the classical substrates of ADAR. Consistent with the use of A-to-I editing as an indicator of bona fide dsRNA, no enrichment was observed for DNA transposons (**Figure 3e**).

To determine if enriched dsRNAs were formed by interactions between separate transcripts (*in trans*), we calculated the proportion of sense or ‘plus’ reads having an overlapping antisense or ‘minus’ read. RNase I-treated samples showed the highest proportion of overlapping sense/antisense reads, with more than 70% of TEs containing reads derived from both strands. This suggests that a substantial fraction of the immunoprecipitated dsRNAs were formed *in trans* (**Figure 3f and Supplementary Figure 4b**). Among TE classes, LINEs and LTRs showed the highest proportions of overlapping reads across all treatments, with the strongest signal observed following RNase I treatment (**Figure 3g**). We observed the same trend when analysing the normalised reads over specific TE examples. These confirmed a clear enrichment of complementary sequences in all dsRNA-immunoprecipitated samples, with the highest number of overlapping sense and antisense reads after RNase I treatment, and an overall lower signal after RNase III digestion (**Figure 3h)**. We hypothesised that the residual signal detected after RNase III treatment is partially due to protection of dsRNA by the J2 antibody. To test this, dsRNA bound to the J2 antibody was recovered after RNase III digestion and subjected to a second RNase III treatment in the absence of antibody. This second digestion resulted in smaller RNA fragment sizes, indicating that the dsRNAs bound to the J2 antibody were partially protected from RNase III cleavage (**Supplementary Figure 4c**).

Together, these analyses indicate that RNase I digestion following J2-based immunoprecipitation enriches for bona fide dsRNAs. This enrichment is supported by higher levels of RNA editing and greater sense/antisense overlap, further confirming the specificity of the identification and characterisation of dsRNA structures in our samples.

### Immunoprecipitated bona fide dsRNAs are enriched in young LINE-1 elements

In order to better understand what TE characteristics are associated with dsRNA formation and RNase I enrichment, we examined additional features of these elements. We first investigated the evolutionary age of the TEs enriched after RNase I treatment using the Kimura divergence of individual copies from the subfamily consensus. This revealed that RNAse I-enriched TEs had significantly lower Kimura divergence values, indicating that they were relatively younger elements (**Figure 4a**). When analysed by TE class, this effect was significant only for LINEs and LTRs, with a much more pronounced effect for LINEs. These findings suggest that younger LINE elements are more likely to form dsRNA and/or generate more complementary, RNase I-resistant dsRNA structures (**Figure 4b**).

**Figure 4.**
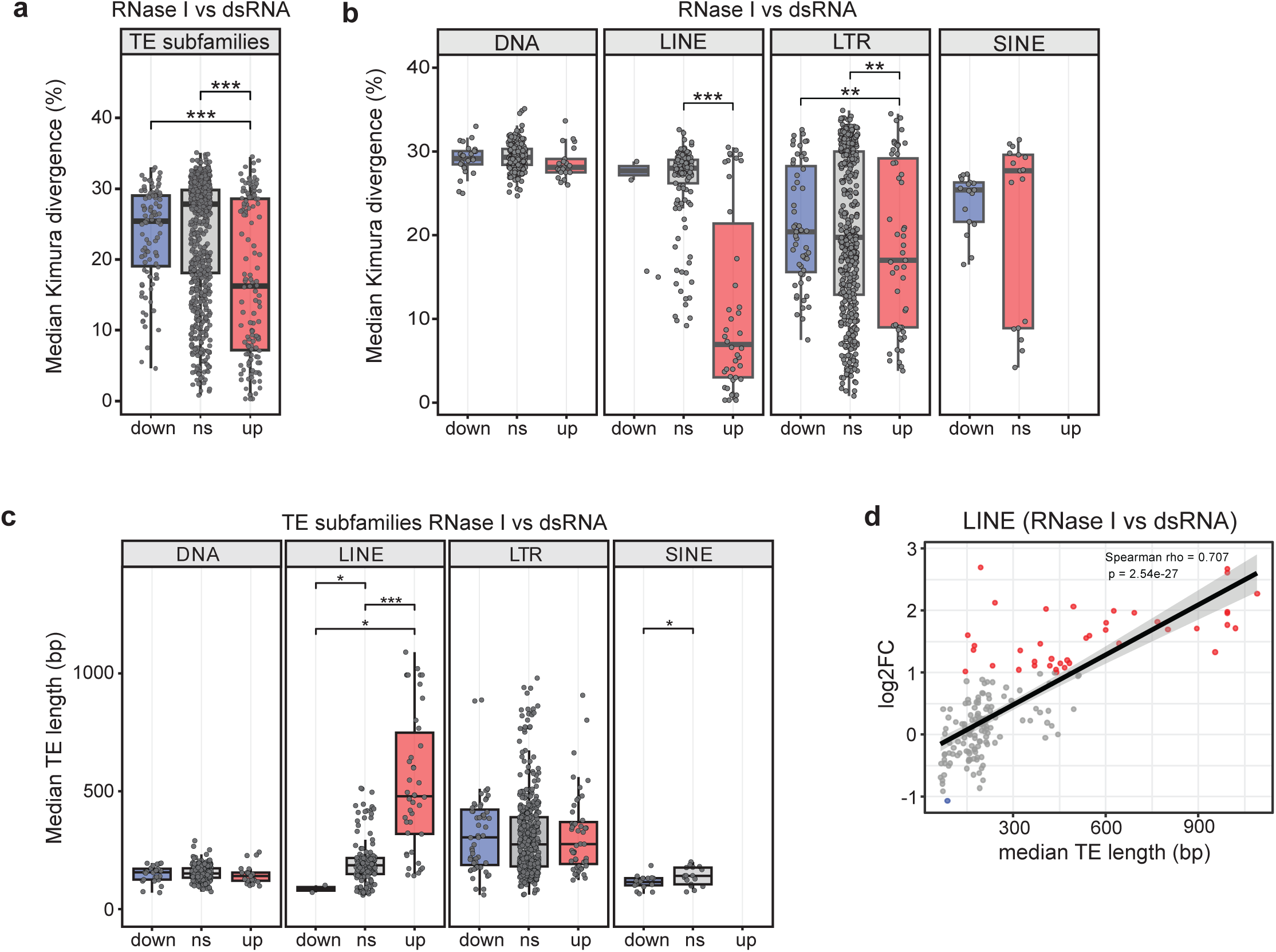
RNase I-enriched TEs are predominantly from young subfamilies. (*A*) Percentage of median Kimura divergence for enriched subfamilies after RNase I treatment versus mock or non-treated samples. Up: upregulated, down: downregulated, ns: non-significant. A Wilcoxon rank-sum test was used to test for significant differences between groups. (*B*) Percentage of median Kimura divergence by TE class. Statistics performed as in (*A*) (*C*) Median TE length in base-pairs (bp) by TE classes enriched upon RNase I treatment vs mock or non-treated. (*D*) Correlation analysis of LINE element-enrichment after RNase I treatment (log2FC, Y-axis) versus length (base-pairs, bp; x-axis). Spearman correlation of rho = 0.707 was significant (p<0.05).

In addition to the evolutionary age, we examined TE length and found that, on average, longer elements were more enriched after RNase I treatment. However, this pattern was specific to LINEs and not observed with the other TE classes (**Figure 4c**). Consistently, LINE enrichment after ssRNA removal showed a significant positive correlation with element length (**Figure 4d**). Collectively, these findings led us to conclude that in ESCs, longer and younger TEs are more likely to form dsRNA. This points to a relationship between the physical properties of TEs and their propensity to form structurally complex RNA molecules, such as dsRNA.

## Discussion

It is becoming increasingly evident that mammalian cells, much like viruses, are capable of producing dsRNA and have evolved multiple mechanisms preventing its accumulation or promote its disruption and degradation. Dysregulation of dsRNA metabolism is now recognised as a contributing factor to human disease, including autoimmune disorders and cancer (de Reuver and Maelfait 2024). Despite these advances, the identification of dsRNA still remains challenging due to the lack of methods that specifically enrich for bona fide dsRNA. Current techniques often require a relatively high-abundance of dsRNA for successful detection, favouring settings with high levels of dsRNA such as infections. The development of methods with an increased sensitivity to bona fide dsRNAs are important in the context of innate immunity as only a small number of dsRNA molecules in a few cells may be required to initiate a cascade of immune reactions.

Intriguingly, ESCs accumulate detectable amounts of endogenous dsRNA, without triggering an antiviral response. Consequently, ESCs have been chosen as a model cell system to refine existing methods for dsRNA identification. Previous methods for dsRNA identification have relied on dsRNA binding proteins or dsRNA-based antibody purification, but these did not include an additional RNase I treatment. Here, we demonstrate that the addition of an RNase I treatment step during the immunoprecipitation enriches for longer and younger LINE and LTR elements. It also results in the enrichment of elements with higher levels of A-to-I editing and increased overlapping sense/antisense reads, suggesting that this extra step enriches for bona fide dsRNAs.

Previous applications of dsRNA-based methodologies have identified both the mitochondria and the nucleus as potential sources of dsRNA in mammalian cells (Sadeq et al. 2021; Chen and Hur 2022; Cottrell et al. 2024). However, we propose that the origin of the dsRNA likely varies depending on the cell type and the nature of the perturbation applied. We do not anticipate that dsRNA identities will be universal across cell types, even they originate from the same genome. For instance, PKR has been shown to interact with mitochondrial RNAs, and the inactivation of the mitochondrial exonuclease Pnpt1 leads to an accumulation of mitochondrial dsRNA in HeLa cells (Kim et al. 2018; Dhir et al. 2018). Nevertheless, in most cases, dsRNA is generated by nuclear transcription, and it is enriched in TEs (Gázquez-Gutiérrez et al. 2021). In agreement, dsRNA-IPs from ESC showed a clear enrichment for TEs but not for mitochondrial RNAs.

One of the best recognised classes of TEs involved in dsRNA formation are SINEs. These elements are the most abundant TEs in mammalian genomes, with approximately 1.4 million copies in the mouse genome and 1.5 million in humans. The successful expansion of SINEs is probably due to being generally well-tolerated even when inserted within genes, as they are small and non-coding. Also, SINE insertions are usually found in the non-coding regions of genes, including introns and 3’UTRs (Lander et al. 2001; Maquat 2020). Their density is so high that a typical human gene contains more than one Alu element, which may be inserted in sense and antisense orientation (inverted-repeat Alus), leading to dsRNA formation. Despite their potential to form dsRNA, SINEs are not predicted to trigger an immune response because they are the major target of ADAR editing activity (Levanon et al. 2024). Despite this, the high abundance of SINEs copies, as well as high number of SINE-overlapping transcripts that are present in cells has made them a prime candidate for dsRNA formation across multiple cell types (Ahmad et al. 2018; Chung et al. 2018; Heinrich et al. 2019; Kim et al. 2014; Mehdipour et al. 2020).

Our data suggest that in addition to SINEs, LTRs and LINEs are also prime candidates for bona fide dsRNA formation in ESCs. We hypothesise that this is likely a direct consequence of the transcriptomic profile of ESCs. During the early stages of development, the expression of other types of full-length elements is temporarily allowed, as the epigenetic marks that silence TEs are remodelled. In ESCs, we observed that bona fide dsRNA is predominantly formed by LINEs and LTRs, as they are proportionally more resistant to RNase I treatment than SINEs. This is consistent with findings where both LINE-1 and LTRs have been involved in dsRNA formation after TE transcription is derepressed, either by inactivating the HUSH complex or by triggering DNA demethylation (Chiappinelli et al. 2015; Roulois et al. 2015; Tunbak et al. 2020). Therefore, we conclude that under conditions where LINEs and LTRs can be expressed, these may also contribute to dsRNA formation. Also, we hypothesise that LINEs and LTRs may be more ‘efficient’ in forming dsRNA, as they tend to be longer than SINEs, and thus the number of potential bases embedded in dsRNA structures is larger.

Interestingly, we observed A-to-I editing across all classes of retrotransposons, including LINEs, LTRs and SINEs. However, SINEs, particularly Alu elements, have been previously identified as the main target for ADAR-mediated editing (Blow et al. 2004; Bazak et al. 2014; Herrmann et al. 2026). We hypothesise that the diversity in editing targets may be linked to the transcriptomic profile of cells. In somatic cells and adult tissues, full-length LINE or LTRs are usually not expressed, and thus they cannot serve as ADAR substrates. Conversely, in ESCs, both LINEs and LTRs retrotransposons are expressed and form dsRNA, thus being able to be edited at levels comparable to those observed in SINEs. The impact of this editing on dsRNA structure may differ among the various TE classes. IR-Alus possibly generate shorter dsRNA stems, up to 300 bp, and contain more unpaired regions that are sensitive to RNase I treatment. In contrast, our findings indicate that LINE elements can form dsRNA with an average length of 500 bp, extending up to 1Kb. Given the strong enrichment of both sense and antisense strands in LINEs, we predict that the dsRNAs formed can be as long as 1Kb. These longer dsRNA segments likely form more stable duplexes which are less affected by editing, while shorter or less stable dsRNA structures, like those formed by SINEs, may be more susceptible to similar levels of editing. Interestingly, the observed levels of editing are high, yet insufficient to disrupt the dsRNA structure enough to prevent recognition by the dsRNA antibody. Although this may not be relevant in the context of ESCs, the extent of editing could still be sufficient to evade cellular immune sensing.

In addition to length, the evolutionary age of TEs appears to be crucial for the formation of bona fide dsRNA. Somatic adult cells rarely express full-length young TEs because they retain the ability to retrotranspose. However, a different scenario unfolds during development. For instance, in mouse genomes LINE and LTR elements, such as LINE- 1 and MusD, are still capable of transposition (Ribet et al. 2004; Richardson et al. 2015). About 3,000 copies from the L1 subfamilies A, Tf and Gf remain active in mice (DeBerardinis and Kazazian 1999; Li et al. 2014). From the perspective of the selfish element, the optimal window for transposition is early development, before germline specification. In this manner, the new transposition event has the potential to be passed on to subsequent generations. Considering that younger elements have had less time to accumulate mutations and be eliminated through post-insertion selection, their transcription in both sense and antisense orientations is more likely to produce perfectly paired dsRNA.

The origin of antisense LINEs and LTRs transcripts remains uncertain. LINE-1 elements contain an antisense RNA-polymerase II promoter, which could potentially lead to dsRNA formation (Li et al. 2014). To our knowledge, no antisense promoters have been identified for LTRs or SINEs (Oomen et al. 2025). However, when mapping dsRNA-seq reads to the consensus sequences of these elements, antisense reads appear not to be confined to a specific region but are detected throughout the element. It is possible that the antisense RNA is co-transcribed from the promoter of its host gene. In humans, LINE-1 insertions within genes tend to be selected in the antisense orientation, possibly to mitigate the harmful effects of sense-oriented copies expressed as part of the host gene (Medstrand et al. 2002; Zhang et al. 2011; Sultana et al. 2019; Flasch et al. 2019). Similarly, in mice, selection appears to have favoured antisense insertions for both LINE and LTR elements, suggesting that expression of the antisense transcript could be driven by the promoter of a neighbouring or the host gene (Zhang et al. 2011). If this model is correct, antisense transcripts that contain LINE and LTR sequences would be constitutively co-expressed embedded in other genes, with dsRNA formation occurring when sense copies are transcribed from their own promoter or as part of the host genes too.

To conclude, we have developed a method to detect bona fide dsRNAs. This approach will be critical for revisiting and confirming the identity of dsRNAs in other contexts, such as autoimmune disorders or cancer.

## Materials and methods

### Cell lines and differentiation

The mouse embryonic stem cell (ESC) line v6.5 was obtained from ThermoFisher (MES1402) and maintained in Dulbecco’s Modified Eagle Medium (DMEMThermoFisher, 41966) supplemented with 15% heat-inactivated foetal calf serum (FCS, ThermoFisher), 1× minimal essential amino acids (ThermoFisher, 11140035), 2 mM L-glutamine, 10³ U/mL leukemia inhibitory factor (Stemcell Technologies, 78056), and 50 µM 2-mercaptoethanol (ThermoFisher, 31350010). ESCs were cultured on 0.1% gelatine-coated plates and dissociated using 0.05% trypsin (ThermoFisher, 25200056). To differentiate, mouse ESCs were grown in ESC medium in the absence of LIF, and supplemented with 10uM of retinoic acid (Sigma-Aldrich, R2625). Cells received fresh medium daily for a minimum of 10 days to ensure successful differentiation.

### dsRNA transfections

Cells were transfected with the dsRNA analogue poly(I:C) (Invivogen, tlrl-pic) using Lipofectamine 2000 (ThermoFisher, 11668027). Transfections were performed in 6 and 24-well format for protein and RNA respectively. At approximately 80% confluency, cells we transfected using 0.5ug/ml poly(I:C) and collected after 2 hours for protein samples and 6 hours for the RNA samples.

### RT-qPCR

Total RNA was extracted using Tri-reagent (MilliporeSigma), and cDNA was synthesised using M-MLV (Promega, M1701) in accordance with the manufacturer’s instructions. qPCR reactions were performed using GoTaq qPCR mastermix (Promega, A6001) using previously published primers (Supplementary file 1) on a QuantStudio 5 (ThermoFisher). Data was analysed using Quantstudio Design & Analysis software. Differences were analysed by single factor ANOVA to calculate significant differences amongst comparisons, followed by a Tukey HSD. *p-val ≤ 0.05, **p-val ≤ 0.01, ***p-val ≤ 0.001.

### Western blots

Cells were washed twice with PBS followed by lysis in RIPA buffer (50 mM TRIS-HCl, pH 7.4, 1% triton X-100, 0.5% Na-deoxycholate, 0.1% SDS, 150 mM NaCl) supplemented with protease (Sigma-Aldrich, 11873580001) and phosphatase inhibitors (Sigma-Aldrich, P7526). Samples were sonicated and centrifugation to shear nucleic acids and clarify the lysates. Protein concentrations were determined using the BCA assay (ThermoFisher, 23225) and equal amounts mixed with LDS sample buffer (ThermoFisher, NP0007) and Sample reducing agent (ThermoFisher, NP0009) and run on 4–12% Bis-Tris precast gels (ThermoFisher). Proteins were transferred to nitrocellulose membrane using wet transfer (mini blot module, ThermoFisher) according to the manufacturer’s recommendations. Membranes were blocked for 1 h at room temperature in TBS-T (0.1% Tween-20) and 5% foetal calf serum before overnight incubation at 4 °C with primary antibody, followed by the appropriate secondary antibody for 1h at RT. Antibodies used were anti-phospho PKR (07-886, Merck), anti-phospho IRF3 (D6O1M, Cell signalling), anti-phospho Eif2a (D9G8, Cell signalling), anti-tubulin (CP06, Calbiochem), anti-GAPDH (CB1001, Merck), anti-mouse 680RD (925-68070, LICORbio), anti-mouse 800CW (926-32210, LICORbio), anti-rabbit 680RD (925-68071, LICORbio), anti-rabbit 800CW (926-32211, LICORbio). Proteins bands were visualised on a Li-cor imaging system (LICORbio). Protein bands were quantified using the Li-cor software and expression levels calculated normalised to tubulin or GAPDH.

### Flow cytometry

Cells were dissociated using 0.05% Trypsin, washed in PBS and resuspended in FACS buffer (PBS with 1% FBS). Cells were pelleted and resuspended in Fixation buffer (BioLegend, 420801,) and incubated for 20 min. at 4°C, washed twice with Intracellular Staining Permeabilisation Wash Buffer (BioLegend, 421002). Cells were washed once with RNase III reaction buffer (NEB, ShortCut,) in the absence of MnCl2 and incubated for 45 minutes at 37°C in 100 µl complete reaction buffer with (2ul) or without RNase III. Cells were washed twice with Intracellular Staining Permeabilisation Wash Buffer and incubated overnight with the anti-dsRNA antibody J2 (English & Scientific Consulting) in Intracellular Staining Permeabilisation Wash Buffer (1:400). After incubation, cells were washed twice with Intracellular Staining Permeabilisation Wash Buffer and incubated with anti-mouse Alexa Fluor 647 (BioLegend, 405322,) for 1 h at room temperature and washed three times with Intracellular Staining Permeabilisation Wash Buffer before finally resuspending the cells in FACS buffer. Cells were analysed using a MACS Quant analyser 10 (Miltenyi), and data were processed using FlowJo software (Treestar).

### Immunofluorescence

Glass coverslips were coated with 0.03 μg/mL laminin (L2020, Sigma Aldrich) for 30 min at room temperature, aspirated and washed with PBS before addition of cells. For immunofluorescence, cells were washed once with PBS and fixed with 4% paraformaldehyde (Sigma Aldrich) for 10 minutes at RT. Cells were washed thrice with PBS, and the cell membrane was stained with 2 μg/mL CF488A-conjugated lectin antibody (29022, biotinum) for 30 min at RT. Cells were washed and permeabilised with 0.2% Triton-X100 (Sigma Aldrich) and incubated in blocking buffer (5% FCS, 0.1% Triton-X100). Cells were incubated overnight at 4°C with J2 antibody (1:300) diluted in blocking buffer. Cells were washed thrice with blocking buffer before incubation with anti-mouse Alexa Fluor 647 diluted in blocking buffer (1:4000) and incubated for 45 minutes at RT. DAPI (Sigma Aldrich) was diluted to 10 μg/mL to stain DNA for 5 minutes at RT. Cells were again washed thrice with blocking buffer before a final wash with PBS prior mounting the coverslips onto microscope slides with ProLong Diamond reagent (P36965, Invitrogen). Image acquisition was performed using a Zeiss Axio Imager Z2. FIJI (https://www.nature.com/articles/nmeth.2019) was used for image analysis.

### DsRNA immunoprecipitation and sequencing

ESCs were detached with 0.05% trypsin and washed twice with PBS. For each IP, 1 15cm plate was used. Cells were lysed in IP buffer (50 mM Tris pH 7.5, 150 mM NaCl, 1 mM EDTA, 1% Triton X-100), supplemented with protease inhibitors (11836170001, Roche), 80U RNase inhibitor (N2518, Promega), 10U DNAse (RQ1, Promega) on ice 30 min followed by centrifugation at 17000xg for 15 minutes at 4°C to clarify the samples. Protein G magnetic beads (50 µl, 10004D, Invitrogen) were washed thrice with IP buffer, and mixed with 5 µg of either anti-dsRNA J2 antibody or IgG2a-κ antibody as a negative control. The conjugated beads were washed thrice with cold IP buffer, and incubated with cell lysates for 4 hours at 4°C. Beads were washed three times with cold IP buffer and incubated with either 100U RNase I (M0243S, NEB) or 2U RNAse III (M0245S, NEB) 100 µl of the manufacturer’s supplied buffers for 20’ at 37°C. Mock-treated samples were resuspended in RNase III buffer without RNase III. After washing three times with IP buffer, RNA was isolated using Trizol LS reagent (T3934, Sigma Aldrich) and 0.3 M sodium acetate, according to the manufacturer’s instructions.

### DsRNA computational analysis

Total RNA quality was assessed on the Agilent 2100 Electrophoresis Bioanalyser Instrument (Agilent, G2939AA). RNA was quantified using the Qubit 2.0 Fluorometer (ThermoFisher, Q32866). DNA contamination was quantified using the Qubit dsDNA HS assay kit and confirmed to be <6% (ThermoFisher, Q32854). Libraries for total RNA sequencing were prepared using the NEBNext Ultra 2 Directional RNA library prep kit (Illumina, E7760) and the NEBNext rRNA Depletion kit (Human/Mouse/Rat) (Illumina, E6310) according to the provided protocol. Sequencing was performed on the NextSeq 2000 platform (Illumina, 20038897) using NextSeq 2000 P3 Reagents (200 Cycles, 2×100bp) (20040559). Paired-end RNA-seq reads were adapter-trimmed using Cutadapt (v5.1) with a minimum length of 50 bp and NextSeq-specific quality trimming (Martin 2011). Reads were aligned to the mouse reference genome (mm39) using STAR (v2.7.11b) with end-to-end alignment, allowing multimapping reads (up to 1000 loci) and a maximum of three mismatches (Dobin et al. 2013). Sorted BAM files were generated and indexed using SAMtools (v1.23.1) (Dobin et al. 2013). Strand-specific coverage tracks were produced with deepTools (v3.5.1) (bamCoverage) using CPM normalization and an effective genome size of 2.6 Gb (Dobin et al. 2013). Sense and antisense signals were intersected using BEDTools (v2.31.0) to quantify strand overlap (Quinlan and Hall 2010). dsRNA levels were quantified using an overlap metric, defined as the proportion of antisense signal overlapping sense transcription (overlap / sense signal). Overlap was calculated as the minimum coverage between strands at each position, weighted by genomic span. Metrics were computed genome-wide and within TE classes using merged RepeatMasker-derived annotations.

Reads were assigned to genomic features using featureCounts (v2.1.1) with multimapping enabled (Liao et al. 2014). Gene-level counts were summarized by transcript biotype, while repeat counts were grouped by TE family and class using custom annotations. Counts were normalized per sample to relative proportions.

Transposable element (TE) and gene counts were quantified using TEcount (TEtranscripts v2.2.3) in multi-mapping mode with strandedness set to reverse, using gene and TE annotations (Jin et al. 2015). Differential expression analysis was performed in R (v4.5.0) using DESeq2 (v1.44.0) (Love et al. 2014). Size factors were estimated from gene counts only and applied to TE counts to avoid bias from repetitive elements. Lowly expressed features (mean count <10) were excluded. Differential expression was assessed using a negative binomial model, and significance was defined as |log2FC| > 1 and adjusted p < 0.05. Variance-stabilizing transformation (VST) was applied for downstream visualization.

RNA editing was assessed using REDItools3 (v3.6) by comparing aligned reads to the reference genome (Picardi and Pesole 2013; Fonzino et al. 2025). Editing indices were calculated across different TE annotations.

## Data availability

All Illumina sequencing datasets have been deposited under the SRA BioProject PRJNA1441232. Raw data can be accessed at the link (https://dataview.ncbi.nlm.nih.gov/object/PRJNA1441232?reviewer=a9n2uphg9ns6ca6opfkb7inglv)

## Acknowledgements

This work was funded by the Leverhulme Trust (RPG-2020-355) and the Wellcome Trust grants (221737/Z/20/Z) and (107665/Z/15/Z) to S.M. The S.R.H group is supported by the Spanish Ministry of Science and Innovation (PID2024-162606NB-100 and CNS2023-145402). K.K. is supported by a Darwin Trust fellowship. We thank Felix Mueller for help with the J2-flow cytometry assay.

## Author contribution

K.K and J.W performed all the wet lab experiments. P.G.M and G.P performed the computational analyses, under the supervision of J.W., S.R.H and S.M. J.W., K.K and P.G.M prepared the figures. J.W. and S.M wrote the manuscript with help of all the authors.

